# Evaluation of Human Norovirus Replication in Salivary Gland Cells

**DOI:** 10.64898/2026.09.29.755260

**Authors:** Jennifer Harlow, Simon Briggs, Emma Pothier, Katherine Pham, Neda Nasheri

## Abstract

Human noroviruses (HuNoVs) are the most common cause of non-bacterial foodborne illnesses worldwide. HuNoVs are notoriously difficult to culture, and current cell culture systems, such as human intestinal enteroids have limitations in efficiency and scalability, which has limited their applications in food safety studies. Recently, it has been shown that transformed salivary gland cell lines (including NS-SV-TT-DC) allow for replication and passaging of norovirus GII.4. Herein, we examined whether we can independently observe viral replication in salivary gland cells and if other strains of human norovirus can replicate in these cells. For this purpose, we first evaluated the replication of several norovirus strains in NS-SV-TT-DC cells by either infecting them with vesicle-cloaked viruses or total viruses in stool filtrates. Low levels of viral replication was observed and vesicle isolation did not improve viral replication. Next, we screened for replication of 50 norovirus samples from 9 genotypes, and 14 isolates demonstrated over a two-fold viral replication. We observed that the initial viral load in the samples does not have a significant effect on replication efficiency. We were also able to successfully passage several isolates. Subsequently, we examined whether interferon inhibitors such as Ruxolitinib, TPCA-1, and BX795 would improve viral replication and observed negligible effects. Finally we tested multiple supplements including MgCl_2_, bile salt, ceramide, and TAK-779. Most supplements had little or no effect on HuNoV replication in NS-SV-TT-DC cells. Bile salt and TAK-779 showed limited, genotype-specific benefits. Altogether these results indicate that salivary gland cells allow limited replication of certain HuNoV isolates.

**Importance:** The lack of robust and accessible cell culture systems continues to hinder human norovirus research. Salivary gland cell lines have recently emerged as a promising alternative to human intestinal enteroids because they are easy to maintain and relatively unexpensive. Here, we provide a large-scale independent evaluation of the NS-SV-TT-DC salivary gland model using 50 clinical isolates from multiple human norovirus genotypes. Although several isolates replicated and could be serially passaged, replication was typically modest. Furthermore, multiple approaches commonly used to enhance viral replication in cell culture had little or no effect on system efficiency. These findings establish realistic expectations for the use of this model and provide a framework for future optimization of human norovirus cultivation systems.

## Introduction

Human norovirus (HuNoV) is a major cause of foodborne illness worldwide and is estimated to cause approximately 233 million cases and 62,000 deaths of acute gastroenteritis per year, as well as 4 million disability-adjusted life-years (DALYs) globally (1). The illness caused by HuNoV is usually acute and resolves without medical intervention; however, chronic or severe infections leading to death could also occur (2). Based on genetic diversity, noroviruses are classified into at least ten genogroups I-X with GI, GII, and GIV infecting humans. Noroviruses are further divided into 49 genotypes and 60 polymerase types (3).

HuNoVs cannot be easily cultivated, therefore they are considered as “non-culturable enteric viruses” (4). Since the ability to test the viral infectivity and propagate the viral stocks *in vitro* has immense benefit to food safety, antiviral and vaccine development, multiple systems have been developed and evaluated for HuNoV replication (5). Stem cell-derived human enteroids (HIEs) have been widely used for replication of HuNoVs from several genotypes for various research purposes (6, 7). However, the HIE system is costly and cumbersome, which has impacted its application in food safety testing (8). Hence there is a need for a much simpler system to reliably cultivate HuNoV extracted from a variety of samples while using less resources. In 2022 Ghosh et al reported that SV-40 transformed salivary gland cell lines NS-SV-TT-DC (ductal) and NS-SV-TT-AC (acinar) support the replication of the HuNoV GII.4 Sydney 2012 strain with an efficiency approximated to those reported for the HIE cultures (9). The advantage of these cell lines is that they grow as a monolayer and could be routinely cultured. For HuNoV replication, positive stool filtrate is inoculated with the cells and is subsequently washed off at 6 hours post-infection (hpi).

Viral replication is assessed at 96hpi by measuring the increase in viral genome copies compared to 6hpi. The authors have shown that NS-SV-TT-DC (ductal) cells were more efficient in supporting the replication of HuNoV GII.4. Since working with this replication model is simpler and requires less resources compared with the HIE system, we sought to independently evaluate its efficiency by examining the replication of multiple HuNoVs from various genotypes and investigate the effects of serial passaging, interferon (IFN) inhibitors and media supplements in improving it.

## Materials and Methods

### Cell culture

Human salivary gland cells, an SV40 transformed adherent ductal cell line, NS-SV-TT-DC, was kindly provided by Dr. Nihal Altan-Bonnet at the National Institutes of Health, Bethesda, MD, USA. The cells were maintained with the Keratinocyte Growth Medium 2 Kit (Cedarlane, Cat # C-20111) and supplemented with 10% FBS.

### HuNoV strains

HuNoV isolates used in this study were either submitted to our reference centre for diagnosis or were provided by Public Health Ontario. Presence of norovirus was confirmed and quantified using the procedures explained previously (10).

### Vesicle isolation from human stool samples

Vesicle isolation was conducted as described by Ghosh et al (Ghosh et al., 2022). Fecal samples were diluted in 50% vol/vol in PBS. Aliquots of 1 mL were centrifuged to clear stool samples of debris at 4 °C as follows: 500 × g for 10 min; 1,000 × g for 10 min; 2,000 × g for 10 min; 4,000 × g for 10 min; 7,000 × g for 10 min 3 times. Stool debris pellets were discarded and the supernatant was transferred to a new microcentrifuge tube. Vesicles were isolated from cleared stool solutions using the MagCapture Exosome Isolation Kit PS (catalogue no. 293-77601; Fujifilm Wako) according to the manufacturer’s instructions.

### Viral infection and passaging

Cells were seeded in 12-well plates at a concentration of 4 x 10^5^ cells per well and incubated until 100% confluent monolayers were formed, approximately 3-4 days. Cells were then inoculated with 10 μl of viral suspensions in 200 µl serum free media (210µl total per well) and incubated for 6 h before washing and replacing with fresh medium. Following a total of 96 h of incubation, cells and supernatant were freeze–thawed at −80°C for a total of three cycles and centrifuged for 10 min at 4°C at 2000 × g. The supernatant was kept for RNA extraction and droplet-digital RT-PCR analysis, and to infect naïve NS-SV-TT-DC cells for further passaging.

### IFN inhibitors and supplement treatments

Ruxolitinib (Cedarlane, Cat # HY-50856), TPCA-1 (Cedarlane, Cat # HY-10074), BX795 (Cedarlane, Cat # HY-10514), and TAK-779 (Cedarlane, Cat # HY-13406) were prepared as 10 mM stocks in dimethyl sulfoxide (DMSO) and used at the indicated concentrations. C2-ceramide (Millipore Sigma, Cat # 860502P) and Glycochenodeoxycholic acid (GCDCA) (Millipore Sigma, Cat # 700266P) were also prepared in DMSO and used at the indicated concentrations.

The products were incubated with the cells at indicated concentrations both during virus inoculation (6h) and the subsequent incubation (90h).

### RNA extraction and quantification

For RNA extraction, 140 µl aliquot of viral suspension were used for the QIAamp Viral RNA Mini Kit (QIAGEN, Mississauga, ON, Canada) according to the manufacturer’s instructions. Viral titers were determined by droplet digital PCR (Bio-Rad, Mississauga, Ontario, Canada) using the probes and primers that were described previously(11).

### Statistical analysis

Statistical analysis was conducted on experiments comparing viral replication following different treatments using GraphPad Prism 9.0 and by applying two-way ANOVA (Dunnett’s multiple comparisons) test.

### Ethics approval

Ethics approval for this study was obtained from Health Canada and Public Health Agency of Canada Research Ethics Board (Protocol number: REB 2023-053H).

## Results

### HuNoV replication and comparison between vesicle extracted and total filtrate HuNoV

The Ghosh et al study showed that vesicle-cloaked HuNoV replicates more efficiently compared to free HuNoV in NS-SV-TT-DC cells and pediatric samples (< 2 years) have a higher replication efficiency (9). For this purpose we initially examined the replication of 3 pediatric and 4 non-pediatric samples in NS-SV-TT-DC cells and assessed whether vesicle extraction could improve viral replication (Table 1). The initial viral load ranged from 6.8 to 9.3 log genome copies per gram stool (gc/g) in pediatric samples and 7.1-10.5 log gc/g in non-pediatric samples. While low levels of viral replication was observed for all samples with total filtrate ranging 0.7-3.0 fold, 3 of the vesicle-extracted samples failed to replicate, meaning that we did not detect any viral RNA at 96hpi. Overall total filtrate viruses replicated more efficiently compared to vesicle cloaked viruses as evidenced by the ratio of 96h/6h titer and the viral titer at 96hpi (Table 1). For this reason, total filtrate viruses were used for further experiments. Moreover, we did not observe any improved replication in pediatric samples compared to non-pediatric samples, as all non-pediatric isolates replicated in NS-SV-TT-DC cells in contrast to 2 out of 3 pediatric isolates (96/6hpi>1) (Table 1).

**Table 1.** Comparison of the replication efficiency in NS-SV-TT-DC between total viruses extracted from stool filtrates or vesicle-cloaked viruses. Data is the average of at least four replicates. NR is no replication.

| Sample ID | Genotype | log gc/g<br>sample titer | 96/6h<br>Ratio<br>filtrate | log 96h titer<br>filtrate | 96/6h<br>Ratio<br>vesicle | log 96h titer<br>vesicle |
| --- | --- | --- | --- | --- | --- | --- |
| BMH-148 | GII.3 (pediatric) | 7.2 | 0.7±0.2 | 2.5±1.9 | 0.7 ± 0.1 | 3.3 ± 2.4 |
| BMH-152 | GII.6 | 7.4 | 2.4± 0.5 | 3.0±2.3 | 0.8± 0.2 | 2.8 ± 2.1 |
| BMH-153 | GII.4 (pediatric) | 9.3 | 1.1±0.2 | 4.5 ± 3.7 | 0.7± 0.2 | 2.6± 2.0 |
| BMH-157 | GII.3 (pediatric) | 6.8 | 1.9±0.4 | 2.6±1.9 | NR | NR |
| BMH-159 | GII.3 | 10.5 | 1.3± 0.2 | 6.2±5.2 | 1.8 ± 0.3 | 4.7 ± 3.9 |
| BMH-167 | GII.3 | 7.2 | 1.3±0.1 | 2.9±1.7 | NR | NR |
| BMH-168 | GII.17 | 7.1 | 3.0±1.5 | 2.4±2.1 | NR | NR |

### Replication of various norovirus isolates in NS-SV-TT-DC

To further explore the ability different HuNoVs isolates in replication in NS-SV-TT-DC cells, we examined 96/6 hpi ratio for 50 isolates from 6 genotypes and 6 P-types. While viral genome could be detected 96hpi for all tested samples, only 14 isolates showed over 2 fold 96/6hpi ratio (Figure 1). The genotype of these 14 samples were GII.17[P17], GII.2[P16], GII.3[P12], GII.4[P16], and GII.4[P31]. The maximum replication reached to 11 fold (1.04 log) and the median replication efficiency was 1.3 fold across the tested samples. Furthermore, we did not observe any correlation between the viral titer in the samples and replication efficiency (R² = 0.0085) (Figure 2).

**Figure 1.**
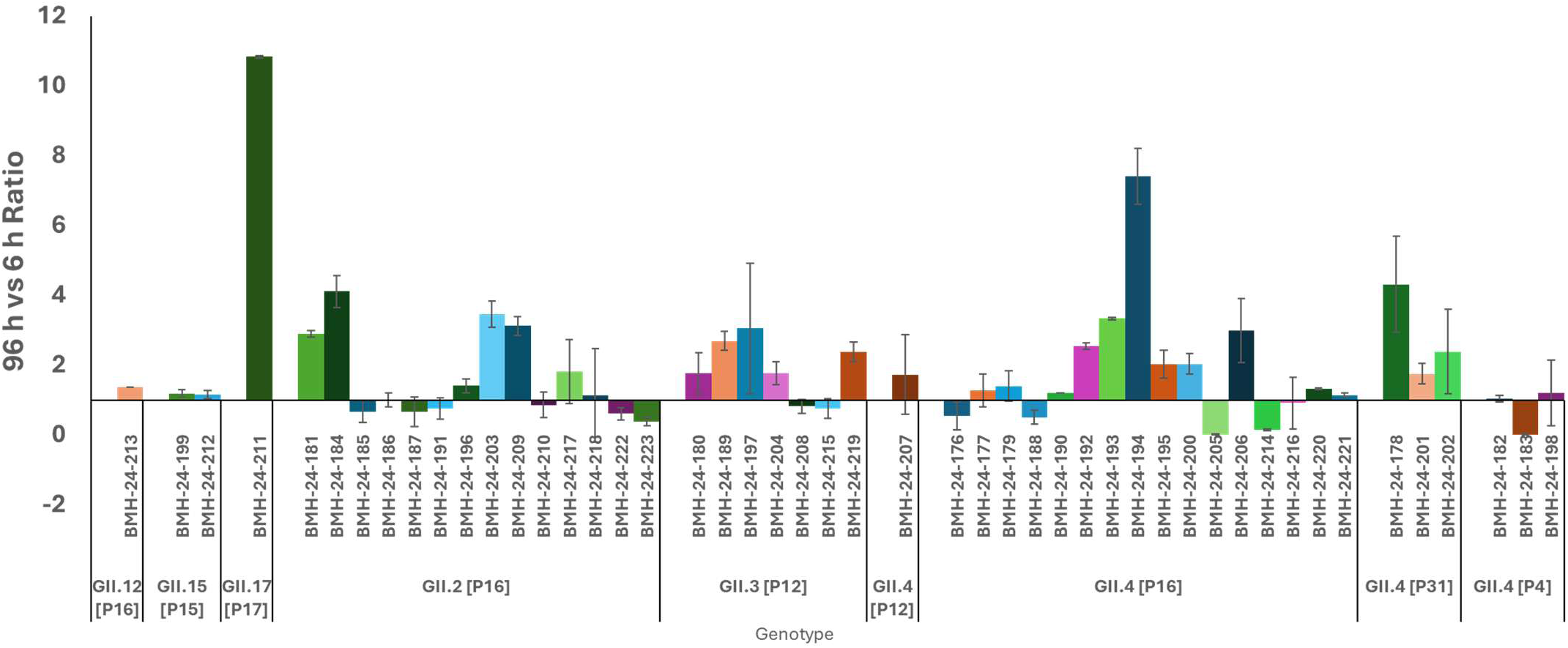
Replication of 50 HuNoV GII samples in NS-SV-TT-DC cells. Over 2-fold 96h/6h ratio was observed for 14 isolates. The data is the average of triplicate experiments and error bars are standard deviation.

**Figure 2.**
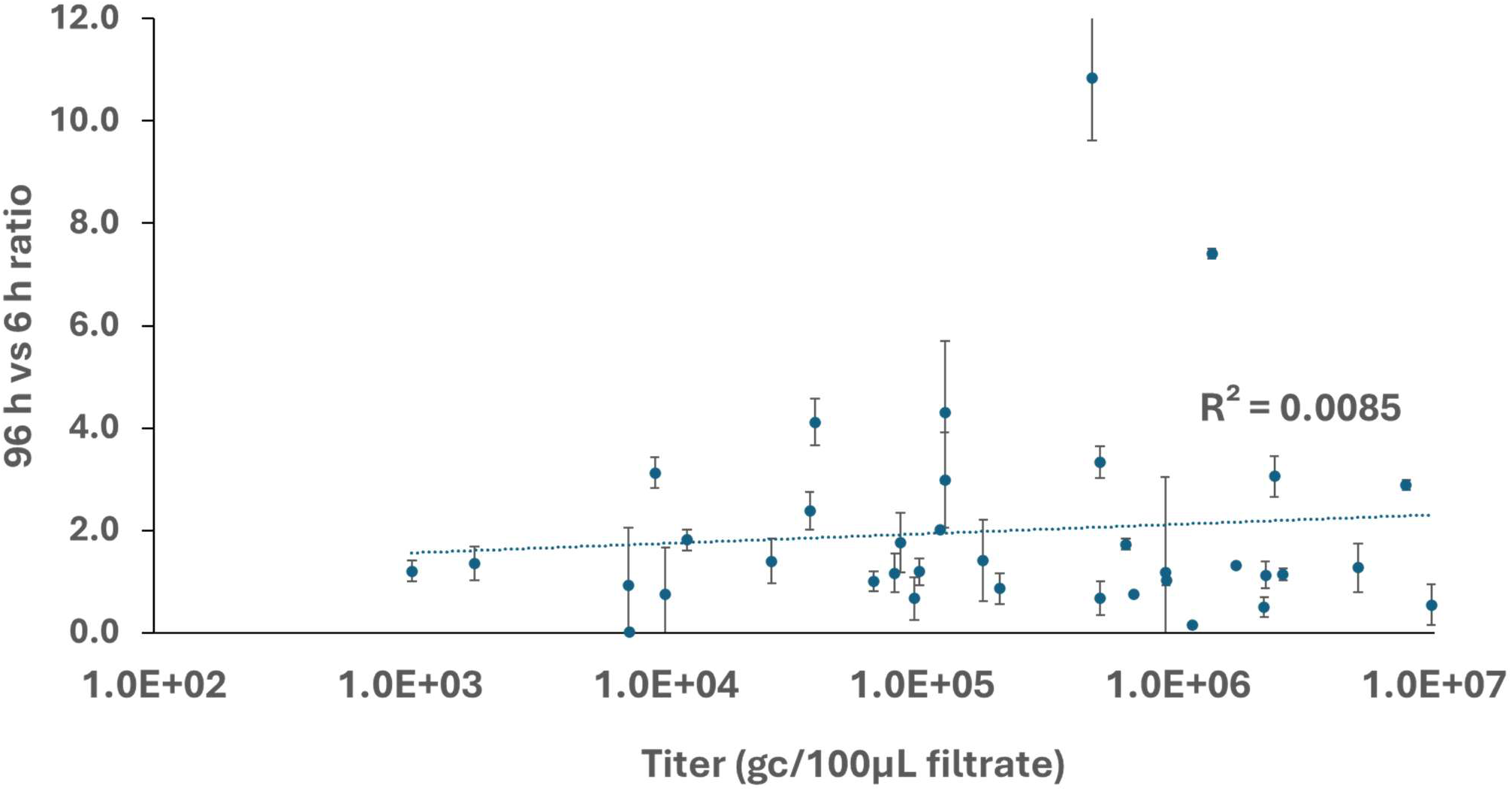
Examining the correlation between sample titer and replication efficiency. We did not find any correlation between the two parameters. Data is the average of triplicate experiments and error bars are standard deviation.

### Passaging the virus in NS-SV-TT-DC

Ghosh et al demonstrated that HuNoV (GII.4 Sydney) could be passaged up to four times in NS-SV-TT-DC cells although the viral titer drops 1 log at P4. We decided to further explore this with other genotypes and also examine if we could adapt the virus to these cells for improved replication efficiency. For this reason we chose several isolates; four GII.4[P16] (177, 179, 193, and 194), one GII.4[P31] (Sydney, 178), one GII.3[P12] (180), one GII.2[P16]) (184) for serial passaging in NS-SV-TT-DC cells. Viral replication was observed for all passaged isolates until P6, and 177 and 178, both GII.4, could be passaged further (P9), however no isolate showed improved replication (Figure 3) as the viral titers did not improve above the P1 levels for any of the tested isolates. Overall these results show that HuNoV could be passaged in NS-SV-TT-DC cells but improved replication was not achieved with the tested isolates.

**Figure 3.**
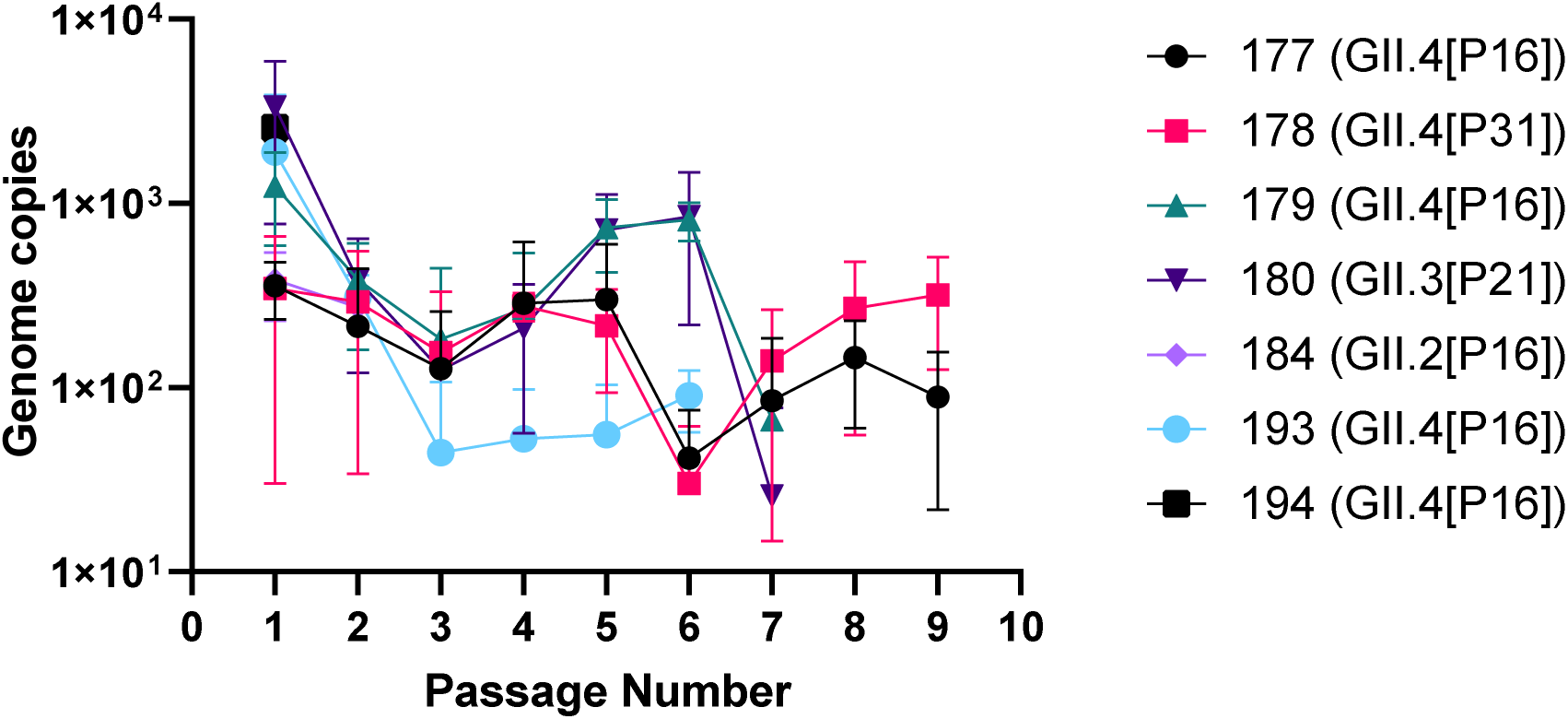
Passaging GII isolates in NS-SV-TT-DC cells. At 96hpi the virus was harvested, quantified and used to infect naïve cells. The data is the average of triplicate experiments and error bars are standard deviation.

### The effect of interferon inhibitors

There are multiple reports that introduction of interferon (IFN) antagonists could enhance viral replication in cell culture (12–14). NS-SV-TT-DC cells are primary human cells, which have been immortalized using SV-40, therefore, it is expected that a strong IFN response is induced following HuNoV infection. For this reason we hypothesized that the inhibition of IFN response would lead to enhanced viral replication. We chose three different IFN antagonists; Ruxolitinib, TPCA-1, and BX-795, that target different pathways and have been shown to enhance viral replication in other systems (14).

We selected 4 HuNoV isolates from different genotypes that showed over 2 fold increase in 96/6 hpi, GII.4[P16] (194 and 203), GII.2[P16] (209), and GII.17[P17] (211). Three different concentrations of Ruxolitinib (0.5-1.5 µM), which targets the Janus kinase1–signal transducer and activator of transcription proteins (JAK1-STAT1) that activates the IFN signaling pathway, were tested and as shown in figure 4A, not only there was no improvement in viral replication compared to the DMSO control, but also it appears that Ruxolitinib had a statistically significant negative effect on viral replication (Figure 4A).

**Figure 4.**
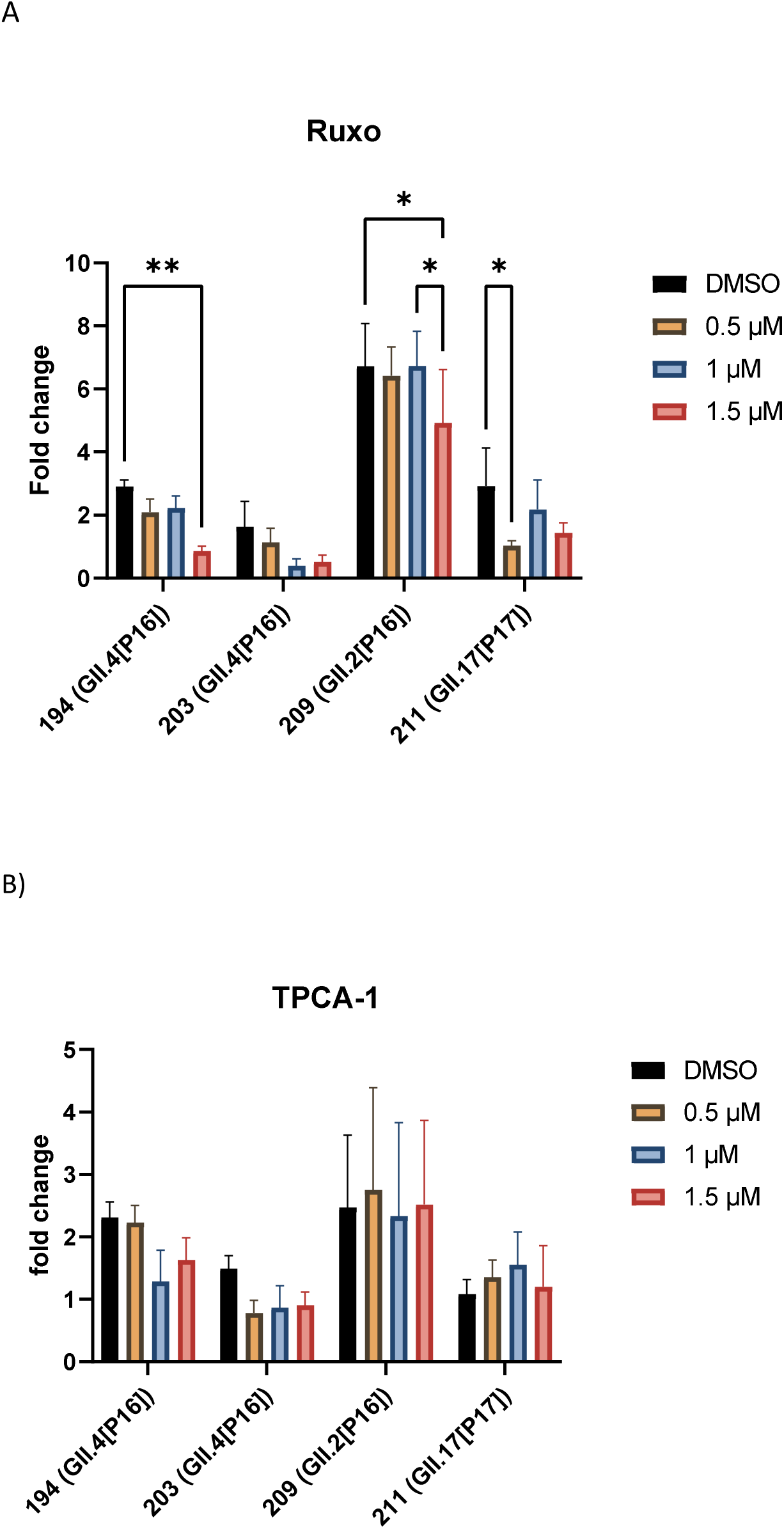
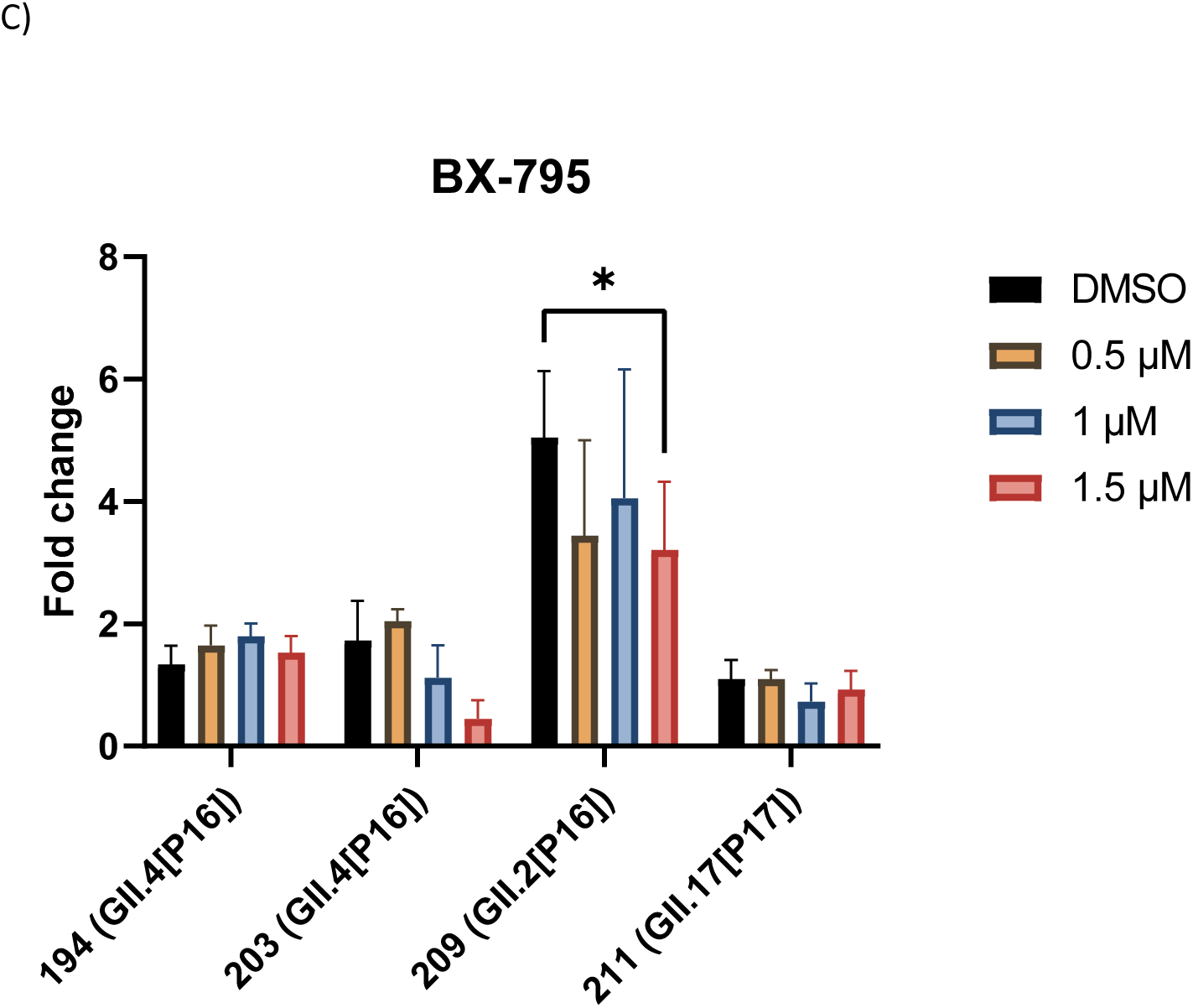
Evaluation of the effect of IFN inhibitors on the replication of HuNoV GII isolates in NS-SV-TT-DC cells. The cells were treated with ruxolitinib (Ruxo) (A), TPCA-1 (B), and BX-795 (C) at the indicated concentrations for the duration of the infection (96h). Data are presented as fold change in viral titer (96/6 hpi) and are the average of three independent experiments and error bars are standard deviation. * P<0.05, ** P<0.01, calculated by 2 way ANOVA.

We next examined the effect of TPCA-1, a dual inhibitor of IKKβ (inhibitor of κB Kinase β) and STAT3 (15), on replication of the four mentioned isolates in NS-SV-TT-DC cells. Studies have demonstrated that TPCA-1 can enhance viral replication at 1µM (16). However, as demonstrated in Figure 4B, presence of TPCA-1 at 0.5-1.5µM did not make any statistically significant improvement to the replication of GII.2 and GII.17 and reduced the replication of GII.4 viruses (Figure 4B).

We also examined whether BX-795, which inhibits TANK-binding kinase 1 (TBK1) that is essential for activation of IRF3 and production of IFN-β, would improve the replication of our candidate GII viruses. We found that addition of BX-795 did not have any positive effect on viral replication and even reduced it in the case of 203 and 209 isolates (Figure 4C). Altogether these data show that IFN inhibitors cannot improve replication of the tested HuNoV GII isolates in NS-SV-TT-DC cells.

### The effect of cell culture supplements

Several reports indicated that supplementing growth medium with MgCl_2_ enhances viral replication in cell culture systems (17, 18). For this purpose, we tested 3 different concentrations (10-30 mM) on the four isolates of HuNoV and as shown in Figure 5A, there was no statistically significant increase in viral replication for GII.4 and GII.17 isolates. However, a mild improvement for the replication of GII.2 isolate (209) at 20mM was observed.

**Figure 5.**
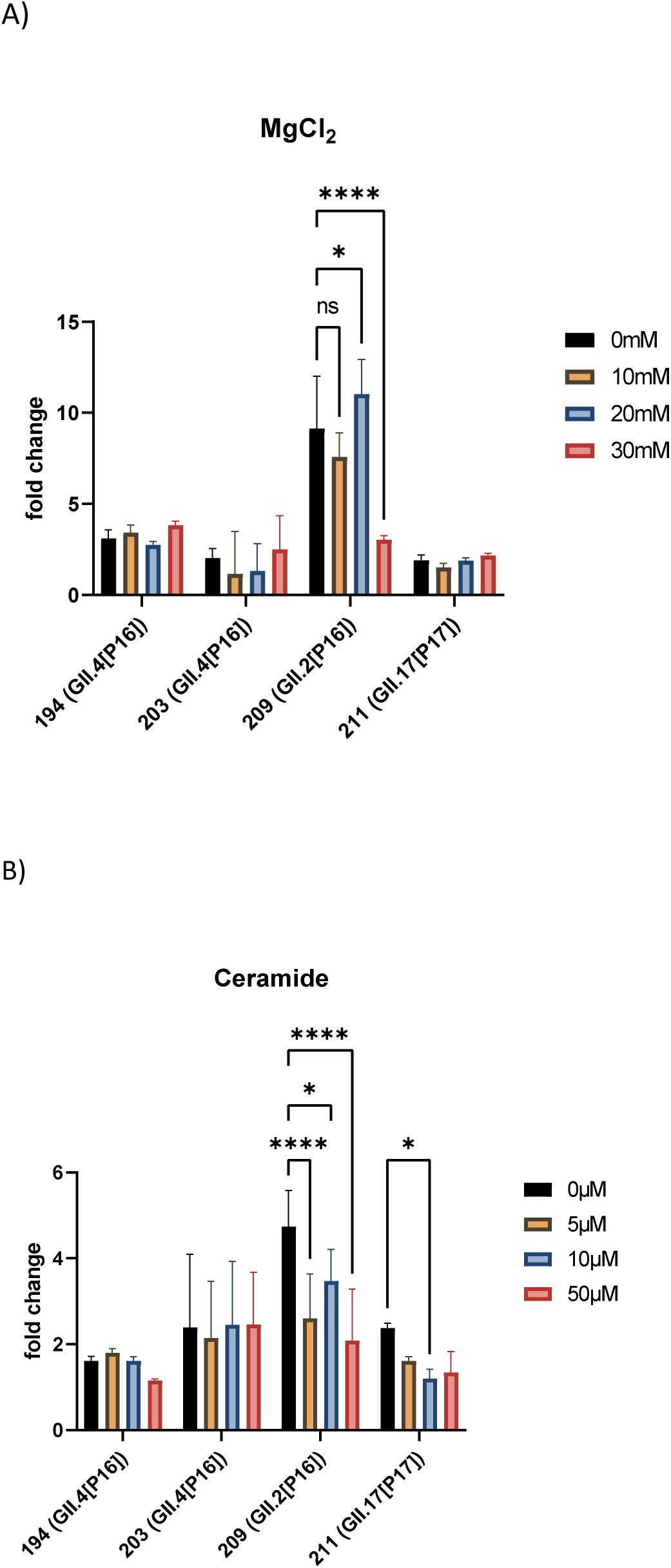
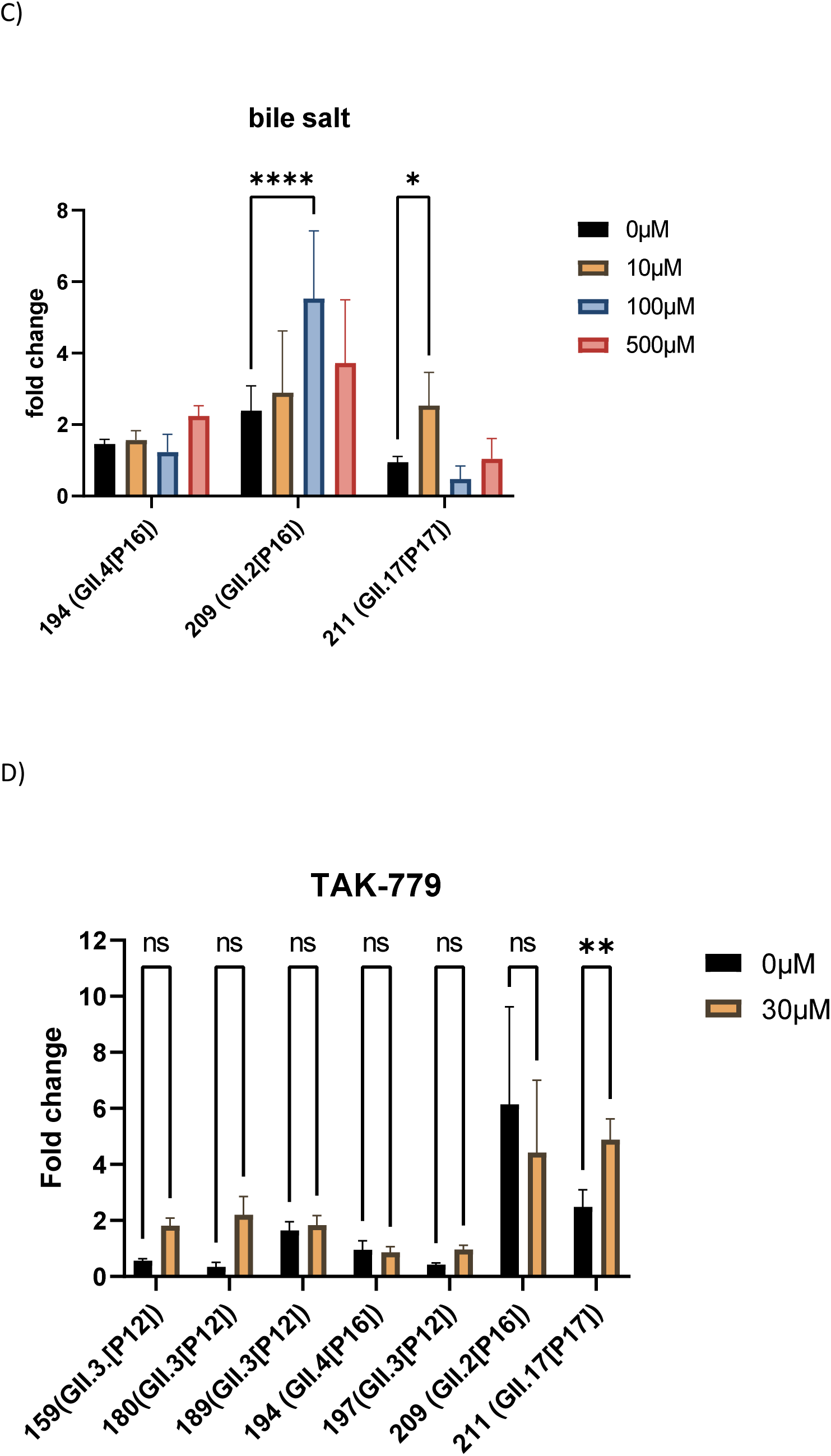
Examining the effect of media supplements on the replication of HuNoV GII isolates in NS-SV-TT-DC cells. The cells were treated with MgCl_2_ (A), C2-Ceramide (B), GCDCA (C), TAK-779 (D), at the indicated concentrations for the duration of the infection (96h). Data are presented as fold change between in viral titer at 96 and 6 hpi and are the average of three independent experiments and error bars are standard deviation. * P<0.05, ** P<0.01, **** P<0.0001, ns non-significant, calculated by 2 way ANOVA.

It has been shown that exogenous C2-ceramide enhances the replication of GII.3 in enteroids (19). To assess whether supplementing the growth media with ceramide would increase the replication of our HuNoV GII isolates in NS-SV-TT-DC cells, we tested 3 different concentrations (5-50µM) on four isolates. As depicted in Figure 5B, supplementing the growth media with ceramide not only did not enhance viral replication, but also slightly reduced it for GII.2 and GII.17 (209 and 211).

Previous studies demonstrated that bile salts, especially glycochenodeoxycholic acid (GCDCA) are essential for the replication of several HuNoV genotypes and enhance the replication of GII.3 in the enteroid system (6, 20). For this reason, we tested a range of concentrations for GCDCA (10-500 µM) on 3 isolates from 3 different genotypes and observed that for GII.2 (209) and GII.17 (211), viral replication is significantly improved in NS-SV-TT-DC cells (Figure 5 C). Encouraged by this result, we tested the replication of more GII.2 and GII.3 isolates in the presence or absence of 100µM GCDCA, however, no significant improvement in replication was observed (Supplementary Figure 1), demonstrating that bile salts are not the limiting factor for viral replication in NS-SV-TT-DC cells.

We next evaluated the effect of TAK-779, a CXCR3/CCR5/CCR2 antagonist that enhanced GII.3 replication in enteroids (21). For this purpose, we examined the replication of four GII.3, one GII.2, one GII.4 and one GII.17 isolates in the presence of 30 µM TAK-779. As shown in Figure 5D, a slight improvement in the replication of all GII.3 isolates was observed but the effect was only statistically significant for the GII.17 isolate (211). No effect was observed for the GII.2 and GII.4 isolates.

We also investigated whether a combination of 20mM MgCl_2_, 100µM GCDCA, and 30µM TAK-779 with or without ceramide would enhance the replication of GII.4 (194) and GII.17 (211). As Shown in Supplementary Figure 2, a combination of the mentioned supplements did not result in a significant improvement in viral replication and addition of ceramide had a negative impact on the replication of the GII.17 (211) isolate.

## Discussion

In an effort to evaluate the replication of HuNoV GII in NS-SV-TT-DC cells and potentially improve on this system, we set out to test a series of genotypes and conditions. Previous studies have reported that HuNoV GII isolates from pediatric patients replicate better in the HIE model compared to isolates from older patients (22, 23). In our pilot screening, all non-pediatric (4/4) samples replicated compared to two out of three pediatric isolates, further, we observed that isolation of vesicle-cloaked viruses negatively affected the replication in NS-SV-TT-DC cells. This could be due to the fact that vesicle isolation drastically reduces the number of viral particles that could initiate infection. Moreover, we observed that high viral titers(gc) does not necessarily lead to higher 96/6 hpi ratio, indicating that many of the RNA fragments quantified by ddRT-PCR cannot establish replication in NS-SV-TT-DC cells.

In this study, the maximum 96/6 ratio was 11 fold (1.04 log) and many isolates failed to replicate. Among the samples that replicated i.e. 96/6>1, the average replication was 2.4 fold, which is closer to 5 fold (0.7 log) increase reported by Kosgei et al (24) than the 3 log increase observed by Ghosh and colleagues (9). Therefore although this system is cost-effective and allows for low levels of viral replication, further strides should be taken to make it into a robust cell culture system for HuNoVs. For this purpose, we conducted a series of experiments to examine whether viral replication could be improved.

Enhanced viral replication following serial passaging has been demonstrated for several viruses such as Zika, influenza, polio, and SARS-CoV-2 (25–27). Herein we intended to examine if HuNoVs could be passaged in NS-SV-TT-DC cells and whether this process would improve viral replication through adaptation and enhanced viral fitness. Although several isolates could be passed a few times, no viral replication was observed past passage 9, and no improvement was observed in viral replication. Altogether, these data demonstrate that passaging is possible in NS-SV-TT-DC cells but this does not necessarily lead to improved viral replication and adaptation. Future works with serial passaging of certain isolates might lead to viral adaptation and improved replication in NS-SV-TT-DC cells.

Since IFN inhibitors have often been used to increase viral growth *in vitro*, we set out to examine whether they could improve viral replication in salivary gland cells. However we did not observe such effect with the tested IFN inhibitors. With Ruxolitinib treatment, we saw significant reduction in viral replication. The antiviral effect of Ruxolitinib has also been reported against HIV-1 and HIV-2 replication (28), and was attributed to the involvement of the JAK-STAT signalling cascade to the pathways that promote viral replication (28). On the other hand, BX-795 works upstream of IFN induction by targeting TBK1, but it also functions as an inhibitor of PDK1, an upstream regulator of the Akt pathway, which is critical for major cellular processes such as growth and metabolism (29) . These off-target effects could be the reason that we saw a decrease in viral replication following treatment with BX-795. The antiviral effect of BX-795 has also been observed for the replication of herpes simplex viruses (30). Lastly, we did not observe a proviral effect with TPCA-1, which directly inhibits NF-κB and STAT3 pathways (15). Importantly the NF-κB pathway also plays a major role in cell metabolism and growth (31), which could be necessary for norovirus replication in NS-SV-TT-DC cells. Altogether these data demonstrate that the employed IFN inhibitors failed to enhance viral replication in NS-SV-TT-DC cells, which could also suggest that HuNoV replication in these cells are more affected by cellular permissiveness than by IFN response. In other words, IFN signalling might not be the limiting factor for the replication of the tested genotypes in these cells.

We have previously shown that replication of hepatitis E virus (HEV) could be improved by 6 fold by addition of 30mM MgCl_2_ to the growth media, however we only observed a slight increase in GII.2 (209) replication at 20mM MgCl_2_ followed by a decrease in replication at 30mM. For the other isolates supplementation of the growth media with MgCl_2_ did not have any significant effect.

The effect of exogenous ceramide treatment on replication efficiency, is virus-dependent (reviewed in (32). While it can enhance the replication of some flaviviruses such as hepatitis C virus and West Nile virus, it reduces the replication of dengue virus, which is also a flavivirus (33). It has also been shown that treatment with exogenous ceramide supresses the replication of influenza A viruses(34), which is consistent with our findings in this study. This suppression of viral replication is more pronounced with GII.2 and GII.17 than GII.4 isolates, which could indicate that ceramide interferes with viral replication cycle possibly by impacting viral assembly.

GCDCA has already been shown to be essential for the replication of GII.3 and GII.17 isolates in the enteroid system, in our study we observed a slight improvement in replication of certain GII.3, GII.2 and GII.17 isolates, but here again it does not seem that GCDCA is the limiting factor for viral replication in NS-SV-TT-DC cells.

TAK-779 has been shown to enhance the replication of GII.3, GII.17 but not GII.4 in the HIE system. Consistently, we observed that the replication of GII.3 and GII.17 isolates was slightly improved in the NS-SV-TT-DC cells, but TAK-779 did not have any effect on the replication of GII.2 and GII.4 isolates. Thus it appears that the enhancing effect of GCDCA and TAK-779 for viral replication in NS-SV-TT-DC cells is genotype dependent.

## Conclusion

Overall the NS-SV-TT-DC cells support the replication of certain HuNoV isolates and the maximum replication that was achieved in this study was about 1 log, however, these cells are not as efficient as the enteroid systems for replication of HuNoVs. This could be due to the lack of key host factors for the replication of HuNoVs in NS-SV-TT-DC cells. Molecular comparison studies could be conducted between the enteroids and salivary gland cells to identify the lacking host factors in salivary gland cells and for improving the replication efficiency of HuNoVs.

## Acknowledgements

This study is financially supported by the Bureau of Microbial and Toxicological Food Safety at Health Canada. The authors would like to thank Dr. Annika Flint and Dr. David Prescott for reviewing the manuscript and providing insightful comments.

## CRediT author statement

JH: methodology, validation, formal analysis, project administration, supervision, review & editing. SB: methodology, formal analysis. EM: methodology, formal analysis. KP: methodology, formal analysis. NN: conceptualization, funding acquisition, software, investigation, writing, visualization.

## Conflict of interest

The authors declare no conflict of interest.

